# Impacts of Morphology and Elasticity on Cancer Cell Deformation in Shear-flows

**DOI:** 10.64898/2026.02.15.703845

**Authors:** Meraj Ahmed, Lahcen Akerkouch, Aaron Vanyo, Amanda Haage, Trung Bao Le

**Affiliations:** Department of Civil, Construction, and Environmental Engineering, North Dakota State University, 1410 14th N, Fargo, 58102, ND, US.; Department of Biomedical Sciences, University of North Dakota, 1301 N Columbia Rd W315, Grand Forks, 58202, ND, US.

**Keywords:** cancer cell migration, dissipative particle dynamics, Fluid-Structure Interaction, metastasis, cancer cell detection

## Abstract

**Purpose:** This work investigates the role of the cancer cell morphology and elasticity on the deformation patterns under shear-flow in a micro-channel.

**Methods:** A novel hybrid continuum-particle framework is developed to simulate cancer-cell dynamics. Cell membrane and nucleus geometries are reconstructed from microscopic images and modeled using Dissipative Particle Dynamics, while the surrounding blood plasma is treated as an incompressible Newtonian fluid. Cell-flow interactions are captured via an immersed boundary method.

**Results:** All cancer-cell models exhibited a rapid deformation response within the first 1-2 ms, followed by morphology- and stiffness-dependent shape evolution. The compact morphologies showed strong recovery, whereas the other models evolved toward folded/lobed states with only intermittent partial recovery during shape transitions. Membrane stiffening dominated elongation and compactness loss, while nuclear stiffening modulated deformation excursions and partial recovery. These shape transitions were accompanied by near-field vortex reorganization and traction localization. Similar to deformation response the net membrane force exhibited a common start-up rise within 0-0.5 ms followed by relaxation. Compact morphologies produce lower and steadier forces. They show minimal stiffness dependence. Deformation-prone morphologies show stronger unsteadiness and clearer stiffness modulation. Cross-sectional velocity and vorticity fields showed a dominant ***x***-directed hydrodynamic imbalance and lateral migration.

**Conclusion:** Our results demonstrate that morphology sets the stiffness modulated deformation patterns which effects the extracellular flow dynamics and traction. In turn, the resulting flow field and traction distribution feed back to influence subsequent deformation and migration. This mechanistic link provides a framework for interpreting circulating tumor cell transport in shear-dominated metastatic environments.

## 1 Introduction

Cancer metastasis, the leading cause of cancer-related mortality [1, 2], is a multi-step process in which malignant cells disseminate from the primary tumor to distant organs [2, 3]. This metastatic cascade starts with the intravasation of tumor cells into the bloodstream, transport through the circulatory or lymphatic systems, and subsequent colonization at distant sites [1, 3]. Once inside the bloodstream, these disseminated cells are termed circulating tumor cells (CTCs) [1, 4]. CTCs are most commonly detected as individual cells[5, 6].

CTCs are increasingly employed as minimally invasive biomarkers for early cancer detection, prognostic stratification, and longitudinal disease monitoring [7]. However, a detailed mechanistic understanding of their behavior in blood flow remains limited. The extensive heterogeneity of CTC phenotypes and their resemblance to white blood cells often limit specificity, compromise purity, and result in CTC loss during clinical investigation [8]. Numerous technologies have been developed to separate CTCs from blood [9], yet their utility is still constrained by complex device designs, elevated costs, and progressive CTC attrition [10]. Due to the complexity of CTC transport, current technologies are limited by modest throughput, suboptimal patient-level specificity, and incomplete validation [11].

Hydrodynamic forces play a critical role in controlling the cell membrane and nucleus during the microvascular transport[12]. Computational approaches has been developed to model cancer cell transport in microvascular flows [13]. Hybrid strategies such as the lattice Boltzmann-immersed boundary (LBM-IBM) models [14] and the immersed finite element models (IFEM) [15] have been employed to examine CTC squeezing, rolling, and adhesion in three-dimensional microvascular geometries [16]. At the mesoscopic scale, the dissipative particle dynamics (DPD) method has become an increasingly popular tool for simulating blood-cell suspensions, in which both the blood plasma and the cell membranes are represented as interacting particles subject to stochastic and hydrodynamic forces [17]. Boundary integral methods (BIM) [18] and smoothed particle hydrodynamics (SPH) [19] have likewise been applied to capture deformable cell migration. However, existing cell models focus on membrane dynamics [1], which often do not resolve the detailed extracellular flow patterns and the fluid-induced forces on the cellular membrane. In addition, most previous numerical studies used idealized morphology such as spheres [20] or capsules [21], an assumption that is not necessarily realistic for in-situ tumor cells, which often exhibit pronounced morphological irregularity and heterogeneous protrusions [22]. More importantly, the elasticity of cancer cells are generally different from one of their normal counterparts [23]. In many malignancies, filamentous actin content is reduced and redistributed toward the cell periphery, leading to altered cytoskeletal organization and lower effective stiffness [23]. Cell elasticity in turn regulates deformability, which plays an essential role in migration through confined micro-environments [24]. For example, cell deformation amplitude has been shown to inversely scales with membrane and nucleus stiffness [25]. In addition, softer cells also display enhanced translation and higher potential for adhesion in curved or low-shear vessels [26].

In this work, we investigate the impact of cancer cell morphology, membrane, and nucleus stiffness on the deformation patterns under shear-flow in a micro-channel. We aim at resolving both the intracellular mechanics and the extracellular flow patterns to monitor the history of cell deformation[27] in flows. We systematically quantify how membrane and nucleus stiffness modulate cell deformability and the hydrodynamic loads experienced by the cancer cell.

## 2 Methodology

### 2.1 Image acquisition and cell morphology

Breast cancer cells (MDA-MB-231) were cultured on glass coverslips overnight in standard growth medium, as shown in Figure 1a. Cells were fixed with 4% paraformaldehyde and stained for the focal adhesion protein paxillin as previously described [28]. Individual cells were then imaged using a Leica Stellaris confocal microscope with an axial step size of 0.5 *µ*m.

**Fig. 1.**
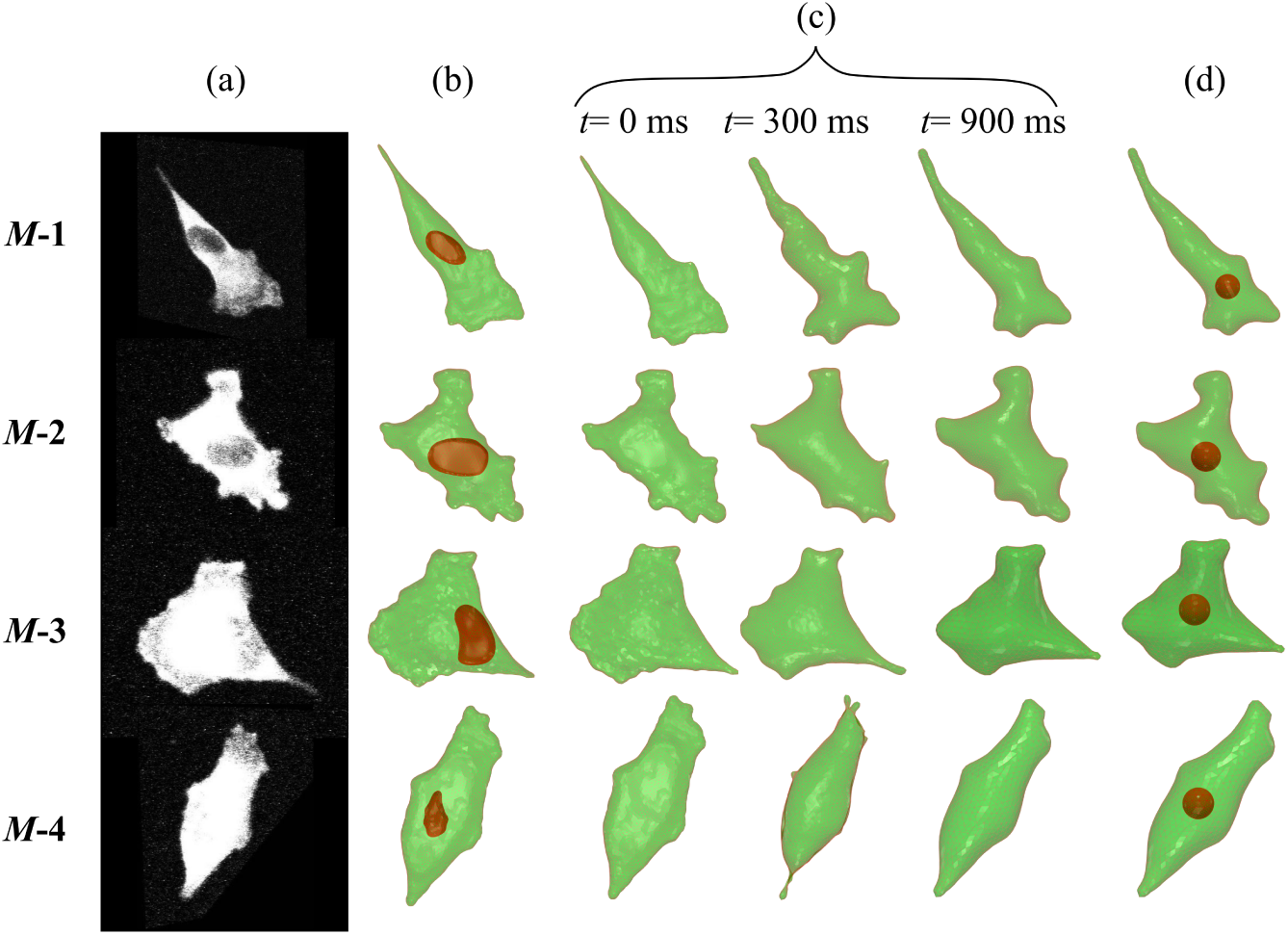
(*a*) Confocal scans of four breast cancer cells type MDA-MB-231 placed on a flat substrate. (*b*) Reconstructed 3D triangulated surface mesh from the confocal scans. (*c*) Snapshot of cell membrane shape during shape relaxation simulation from the original state at *t* = 0 ms to the stress-free equilibrium state at *t* = 900 ms. (*d*) The relaxed cell membranes including the spherical nucleus.

Three-dimensional cancer cell geometries were reconstructed from the confocal image stacks as shown in Figure 1a. These image stacks were processed in the opensource software 3D Slicer to segment the cell membrane by intensity thresholding and generate an initial 3D surface (Figure 1b). The resulting surface meshes were imported into the commercial software Meshmixer, where the triangular mesh was smoothed and remeshed to control resolution while preserving the experimentally observed shape (Figure 1b). In total, there are four cell models: *M* −1, *M* −2, *M* −3, and *M* −4 as shown in Figure 1.

### 2.2 Mechanical model of cancer cell

The geometries of the cancer cell membrane and nucleus membrane were discretized using triangulated surface meshes, as illustrated in Figure 2a. The elasticity of both the membrane and the nucleus was represented by a network of nonlinear springs, in which each edge models the dynamics of spectrin-like links [27]. At each vertex *i*, the dynamics of the cell membrane and nucleus membrane were obtained from the nodal force **F***_i_*, which is derived from the local Helmholtz free energy *V_i_* via

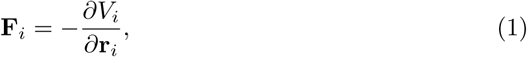

**Fig. 2.**
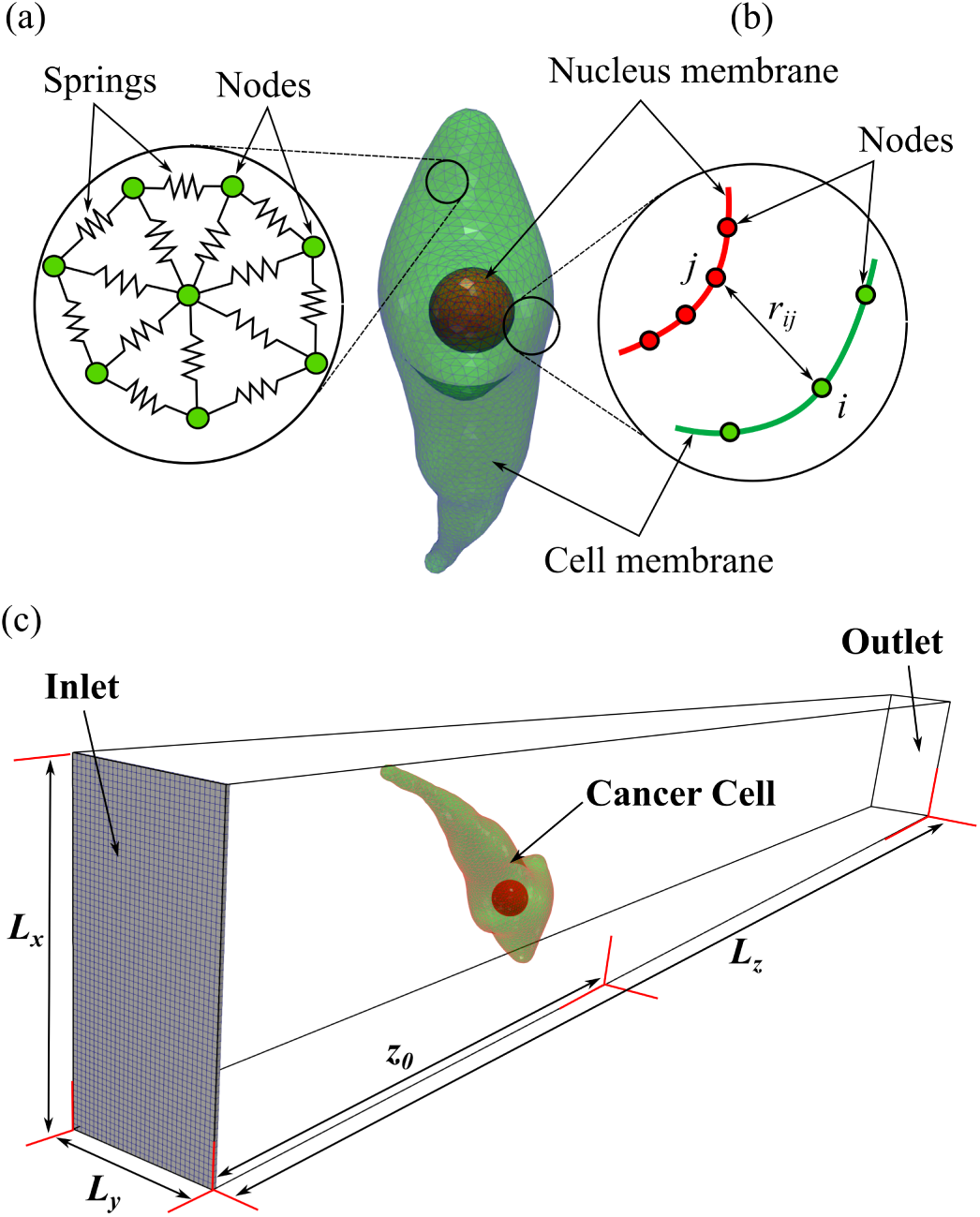
(a) Schematic representation of the cancer cell membrane and nucleus membranes discretized into interconnected nodes and springs in the Dissipative Particle Dynamics (DPD) framework. The membranes are modeled as triangulated surfaces where each node is connected to its neighbors through elastic springs, representing the membrane’s structural stiffness and viscoelastic behavior. (b) The nodal arrangement on both membranes and the separation *r_ij_* between the nodes *i* and *j*. (c) Computational domain for Fluid-Structure Interaction simulation of the cancer cell. The dimensions of the rectangular channel in *x*, *y*, and *z* directions are *L_x_ × L_y_ × L_z_*, see Table 3. The inlet plane of the structured domain is shown in blue to illustrate the computational mesh. The cancer cell is placed at an axial distance *z*_0_ = 90 *µ*m from the inlet plane.

**Table 1.** Physical and model parameters used to characterize the mechanical properties of the cancer cell membrane and nucleus. Here, *D*_0_ is the initial physical diameter, *µ*_0_ the surface shear modulus, *Y* the physical Young’s modulus, and *k_b_* the bending rigidity. *A*_0_ and *V*_0_ denote the initial total surface area and volume, respectively, and *η* is the viscosity of the lipid bilayer. The constraint coefficients *k_a_*, *k_d_*, and *k_v_* correspond to global area, local area, and volume conservation. 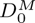, *Y^M^*, and *η^M^* are the corresponding model (dimensionless) quantities used in the numerical formulation.

| Parameter | Membrane | Nucleus | Source |
| --- | --- | --- | --- |
| Initial diameter, $D_0$ [m] | $20 \times 10^{-6}$ | $6.66 \times 10^{-6}$ | Present study |
| Shear modulus, $\mu_0$ [N/m] | $6.3 \times 10^{-6}$ | $6.3 \times 10^{-5}$ | [57] |
| Young’s modulus, $Y$ [N/m] | $2.52 \times 10^{-5}$ | $2.52 \times 10^{-4}$ | Present study |
| Bending rigidity, $k_b$ [J] | $3.6 \times 10^{-18}$ | $5.54 \times 10^{-19}$ | [45, 58] |
| Lipid bilayer viscosity, $\eta$ [Pa · s] | $5.1 \times 10^{-2}$ | $5.1 \times 10^{-2}$ | [59] |
| Boltzmann’s constant, $k_B$ [J/K] | $1.38065 \times 10^{-23}$ | $1.38065 \times 10^{-23}$ | Present study |
| Temperature, $T$ [K] | 298.15 | 298.15 | Present study |
| Global area constraint, $k_a$ [J/m<sup>2</sup>] | $6.17 \times 10^{-5}$ | $1.84 \times 10^{-4}$ | [59] |
| Local area constraint, $k_d$ [J/m<sup>2</sup>] | $1.26 \times 10^{-5}$ | $3.78 \times 10^{-5}$ | [59] |
| Volume constraint, $k_v$ [J/m<sup>3</sup>] | 836.18 | 7525.7 | [59] |
| Model diameter, $D_0^M$ [model units] | 20.07 | 6.77 | [60] |
| Model Young’s modulus, $Y^M$ [model units] | 400 | 400 | [60] |
| Model lipid bilayer viscosity, $\eta^M$ [model units] | 20.4 | 20.4 | [60] |

**Table 2.** Coarse-grained model parameters for the four cancer cell geometries shown in Figure 1. The parameters *l*_0_, *l*_max_, *p*, *k_p_*, and *θ*_0_ are defined in section 2. Nuclear parameters from Model 1 were used for all cancer cell models.

| Coarse-graining | Model 1 |  | Model 2 | Model 3 | Model 4 |
| --- | --- | --- | --- | --- | --- |
|  | Membrane | Nucleus | Membrane | Membrane | Membrane |
| $N_v$ | 2265 | 567 | 1181 | 1080 | 1144 |
| $l_0 \times 10^{-7}$ [m] | 8.02 | 5.29 | 11.1 | 11.6 | 11.2 |
| $l_{\max} \times 10^{-6}$ [m] | 1.76 | 1.16 | 2.44 | 2.55 | 2.48 |
| $p \times 10^{-10}$ [m] | 13.8 | 2.09 | 9.98 | 9.54 | 9.82 |
| $k_p \times 10^{-24}$ [N m<sup>2</sup>] | 1.98 | 5.72 | 5.28 | 6.04 | 5.54 |
| $\theta_0$ [°] | 3.24 | 6.43 | 4.49 | 4.70 | 4.56 |

**Table 3.**
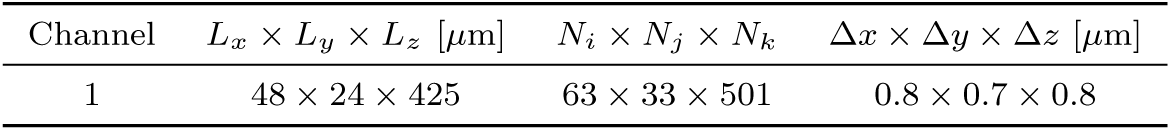
Channel geometry and associated computational grids used to simulate cancer cell dynamics in flow. The channel has a rectangular cross-section and is used for the cancer cell simulations presented in this study. Here, *L_x_*, *L_y_*, and *L_z_* denote the channel dimensions in the axial, spanwise, and wall-normal directions, respectively. *N_i_*, *N_j_*, and *N_k_* are the corresponding numbers of grid points in the *x*, *y*, and *z* directions, and Δ*x*, Δ*y*, and Δ*z* are the associated grid spacings.

where **r***_i_* denotes the position vector of vertex *i*.

The physical properties of the lipid bilayer in the cancer cell membrane and nucleus membrane were modeled using a Helmholtz free energy functional *V* ({**r***_i_*}) that includes contributions from (a) in-plane stretching, (b) bending resistance, and (c) area and volume conservation, such that

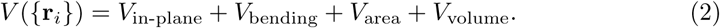

The in-plane free energy *V*_in-plane_ was modeled using a nonlinear Wormlike Chain-Power (WLC-POW) spring formulation:

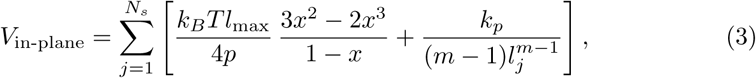

where *N_s_* is the total number of links in the triangulated mesh, and *x* = *l_j_/l*_max_ measures the deformation of spring *j*. Here, *l_j_* is the instantaneous length of spring *j*, *l*_max_ is the maximum allowable link length, *p* is the persistence length, *k_B_*is Boltzmann’s constant, and *T* is the absolute temperature. The parameter *k_p_* is the POW force coefficient, and in this study the exponent was set to *m* = 2 following Fedosov et al. [29].

The bending energy *V*_bending_, which accounts for the resistance of the membrane to bending, was defined as

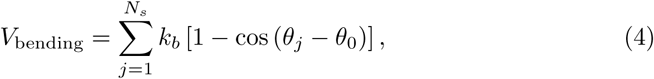

where *k_b_* is the bending rigidity, *θ*_0_ is the spontaneous angle, and *θ_j_* is the instantaneous angle between the normal vectors of two adjacent triangles sharing edge (link) *j*.

The area and volume conservation terms enforce the effective incompressibility of the lipid bilayer and the enclosed cytosol, respectively, and were modeled as

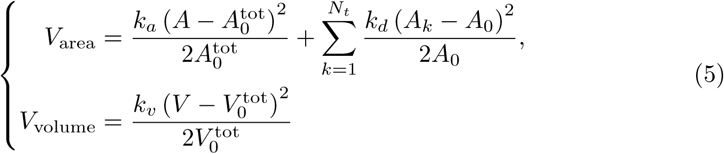

where *N_t_*is the total number of triangular elements. The coefficients *k_a_*, *k_d_*, and *k_v_* are the global area, local area, and volume constraint parameters, respectively. *A_k_* and *A*_0_ denote the instantaneous area of the *k*th triangle and the initial average area per element. 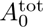 and 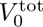 are the equilibrium total area and volume of the cell, and *A* and *V* are the instantaneous total area and volume. The detailed numerical procedure used to evaluate *A* and *V* from the discretized surface was reported in our previous work [27].

#### 2.2.1 Interaction between cell membrane and nucleus: cytoskeleton and cytoplasm model

The physical interaction between the cancer cell membrane and the nucleus is constrained by the intervening cytoskeleton and cytoplasm. To prevent unphysical overlap and to maintain a finite separation between these structures, we introduced a short-range repulsive potential *V*_CSK_ following Ujihara *et al.* [30]:

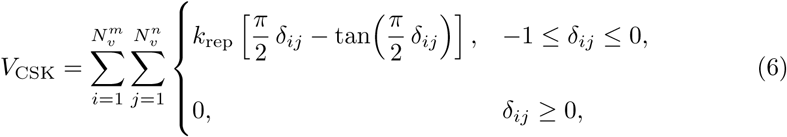

where 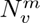 and 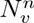 denote the total number of vertices on the cell membrane and nucleus membrane meshes, respectively, and *k*_rep_ is the repulsive stiffness coefficient. The nondimensional gap parameter is defined as *δ_ij_* = (*r_ij_* − *r*_0_)*/r*_0_, where *r_ij_* is the instantaneous distance between the *i*th membrane vertex and the *j*th nuclear vertex, and *r*_0_ is the natural separation corresponding to the difference in the cell membrane and nucleus membrane radii at equilibrium. A schematic of the repulsive interactions between the cell membrane and the nucleus membrane is shown in Figure 2(b).

Equation (1) was used to evaluate the nodal forces associated with each contribution to the Helmholtz free energy *V* in Equations (3)-(5) [27]. Accordingly, for both the cell membrane and the nucleus membrane, the total internal force at vertex *i* was obtained by superposing the individual force contributions,

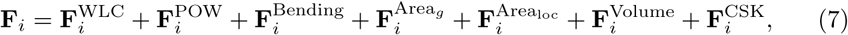

where the superscripts denote the force components arising from the WLC and POW in-plane springs, bending resistance, global and local area constraints, volume conservation, and the cytoskeleton-mediated repulsion, respectively.

#### 2.2.2 Cell Membrane and nucleus membrane viscosity

To account for the viscous response of the cancer cell membrane and nucleus membrane, we incorporated dissipative 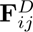 and random 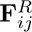 forces by augmenting the spring formulation, following the general framework of the fluid particle model [31]. The forces 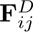 and 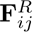 are evaluated for each spring connecting a pair of vertices *i, j* ∈ [1*, …, N_v_*], as given in Equation (8), where *N_v_* denotes either 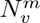 or 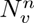 depending on whether the spring belongs to the cell membrane or the nucleus membrane mesh. The relative position and velocity between particles *i* and *j* are defined as **r***_ij_* = **r***_i_* − **r***_j_* and **v***_ij_* = **v***_i_* − **v***_j_*, respectively, with *r_ij_* = |**r***_ij_*| and the corresponding unit vector **r̂***_ij_* = **r***_ij_/r_ij_*.

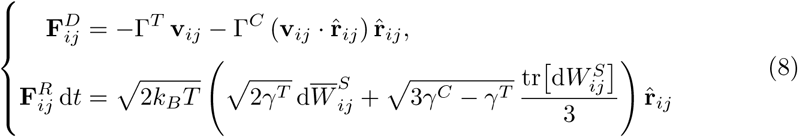

Here, d*t* is the physical time step, tr(d**W***_ij_*) is the trace of the random matrix of independent Wiener increments d**W***_ij_*, and 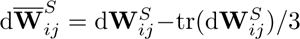 is the traceless symmetric part. The coefficients Γ*^T^* and Γ*^C^* are dissipative parameters, where the superscripts *T* and *C* denote translational and central components, respectively. In this formulation, Γ*^T^* accounts for the dominant contribution to the membrane viscosity, whereas Γ*^C^* provides a smaller correction. Following Fedosov et al. [29], Γ*^C^* is assumed to be one-third of Γ*^T^*, and both parameters are related to the effective surface viscosity of the cell membrane and nucleus membrane, *η*, through

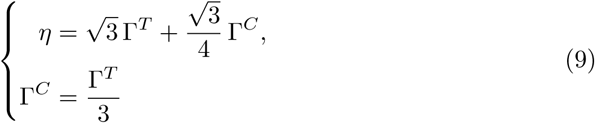

Substituting these relations into Equation (8), the dissipative and random forces between cell membrane and nucleus membrane mesh vertices can be written in terms of the physical viscosity *η* as

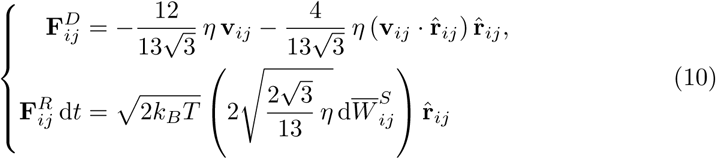

#### 2.2.3 Cell membrane and nucleus membrane elastic properties

The elastic properties of the cell membrane and nucleus membrane, represented as triangulated spring networks, were calibrated using the linear analysis of a twodimensional sheet of springs arranged in equilateral triangles [32]. Building on this framework, Fedosov et al. [29] extended the analysis to a regular hexagonal network endowed with the above energy contributions, thereby establishing explicit relationships between the macroscopic elastic moduli (shear, area-compression, and Young’s moduli) and the underlying model parameters. For the WLC-POW nonlinear spring model, the corresponding linear shear modulus *µ*_0_ is given by

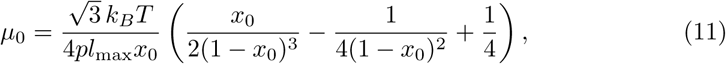

where *l*_0_ is the equilibrium spring length, *x*_0_ = *l*_0_*/l*_max_, *p* is the persistence length, *k_B_* is Boltzmann’s constant, and *T* is the absolute temperature. The linear area-compression modulus *K* and Young’s modulus *Y* of the network are then given by *K* = 2*µ*_0_ + *k_a_* + *k_d_* and *Y* = 4*Kµ*_0_*/*(*K* + *µ*_0_), respectively. Under the incompressibility assumption *µ*_0_ ≪ *k_a_* + *k_d_*, this expression simplifies to *Y* ≈ 4*µ*_0_.

In addition, the model bending coefficient *k_b_* was related to the macroscopic bending rigidity *k_c_*of a spherical membrane in the Helfrich model through *k_b_* = 2*k_c_/*√3 [33].

### 2.3 Fluid-Structure Interaction coupling

The blood plasma was modeled as an incompressible Newtonian fluid governed by the three-dimensional unsteady incompressible Navier-Stokes equations, with density *ρ* and kinematic viscosity *ν* = *µ*_plasma_*/ρ*. In Cartesian tensor notation (with *i* = 1, 2, 3 and Einstein summation over repeated indices), the continuity and momentum equations read

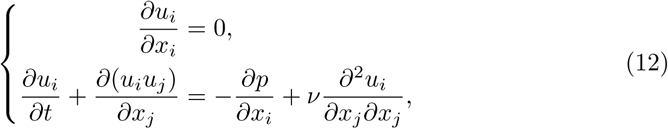

where *u_i_* is the *i*th component of the velocity vector **u**, *x_i_* is the *i*th spatial coordinate, *t* is time, and *p* denotes the pressure scaled by *ρ*. The characteristic velocity scale is taken as *U*_0_, and the reference length scale is set to *L_s_*= 1 *µ*m for all cases.

The fluid equations were solved using a sharp-interface curvilinear immersed boundary (CURVIB) method in a background curvilinear grid that embeds the deformable cell model [34]. The CURVIB framework employed here has been extensively validated for a range of fluid-structure interaction (FSI) problems in biological flows [35–37], therefore, only a brief overview of the coupling strategy is provided. The cellular structure is used to resolve membrane deformation under flow. The FSI procedure couples the motion of the cancer cell membrane to the surrounding plasma via an immersed boundary formulation, in which membrane forces are transferred to the fluid and the local fluid velocity is interpolated back to the membrane nodes. Detailed descriptions of the CURVIB implementation and the FSI algorithm can be found in our previous work [27, 38].

### 2.4 Morphological Analysis

Cell-shape transitions during transport in the microchannel were quantified using sphericity *S* and aspect ratio *AR*. Sphericity, which measures compactness follows Wadell’s classical definition [39],

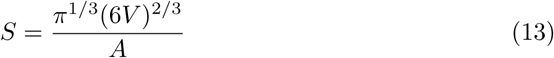

where *V* and *A* are the instantaneous cell volume and surface area, a perfect sphere yields *S* = 1, and lower values indicate reduced compactness. Aspect ratio [40], which quantifies elongation is defined as

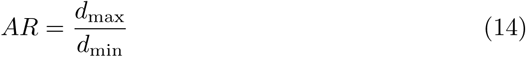

where *d*_max_ and *d*_min_ are the maximum and minimum cell diameters estimated from a bounding box enclosing the cell at each time instant. We propose a new procedure to classify cell morphology during its deformation using two standard indices defined above [41]: i)sphericity (*S*, Eq. 13) and (ii) aspect ratio (*AR*, Eq. 14). Using this approach, cell morphology can be classified into separate groups, which partitions the (*S, AR*) space into five shape categories: (1) round/compact (*R*), (2) amoeboid-ellipsoidal (*AE*), (3) lobed (*DL*), (4) elongated (*EL*), and (5) streamer (*SC*). The physical interpretation of each shape class and other details are given in Table 5.

### 2.5 Computational Setup

The cancer-cell membrane and nuclear envelope are lipid bilayers that resist stretching and bending and therefore store elastic energy when deformed. In the absence of external loads, the system relaxes toward a minimum-energy state. In our model, this relaxed state corresponds to a triangulated surface whose elements are close to equilateral triangles. Because experimentally reconstructed geometries cannot generally be represented using only equilateral triangles, they are not initially in mechanical equilibrium. To obtain relaxed initial shapes, the scanned nuclei were temporarily removed and each cancer-cell model was evolved for 900 ms under force-free conditions (shape-relaxation simulation). The snapshots of cell shapes during shape-relaxation at *t* = 0 ms, 300 ms, and 900 ms are shown in Fig. 1c. The nucleus, which is substantially stiffer than the surrounding cytoplasm, was then reintroduced as a sphere and placed inside the relaxed cell volume (Fig. 1d). These relaxed cell-nucleus configurations were used as initial conditions for the FSI simulations. To examine the role of membrane and nuclear stiffness, each geometry was simulated using the three stiffness combinations listed in Table 4, representing progressively stiffer cell and nuclear envelopes and motivated by cancer-cell softening during metastasis. The computational domain is a rectangular channel containing a single breast cancer cell (Fig. 2c), with dimensions *L_x_* (width), *L_y_* (height), and *L_z_* (length), discretized on a structured grid of size *N_i_* × *N_j_* × *N_k_* with spacings Δ*x*, Δ*y*, and Δ*z*. The reference configuration uses 63 × 33 × 501 cells in *x*, *y*, and *z*. Channel geometries for all cases are summarized in Table 3. Blood plasma viscosity is set to *ν* = *µ*_plasma_*/ρ* = 1.2 × 10*^−^*^6^ m^2^*/*s, and the bulk velocity is *U* = 1 mm*/*s. A linear inlet ramp is applied over *t*_ramp_ = 0.5 ms, during which *U* (*t*) increases from zero to *U*, and is then held constant for the remainder of the simulation. The four cell geometries (*M* −1-*M* −4) were placed in the same channel (Channel-1 in Table 3), at *t* = 0, the cell centroid is located a distance *z*_0_ from the inlet. The effects of membrane and nucleus stiffness were examined by varying the Young’s moduli of the membrane (*Y_m_*) and nucleus (*Y_n_*) from 1.26 × 10*^−^*^5^ to 2.52 × 10*^−^*^5^ N*/*m (i.e., *Y_m_*_1_ to *Y_m_*_2_ and *Y_n_*_1_ to *Y_n_*_2_). Combining four geometries with three stiffness combinations yields 12 simulation cases (Table 4). A physical time step size of 1 *µ*s was used to simulate each case. All simulations were performed on a high-performance computing cluster at North Dakota State University (https://www.ndsu.edu/ccast) using 64 CPUs and 256 GB memory.

**Table 4.**
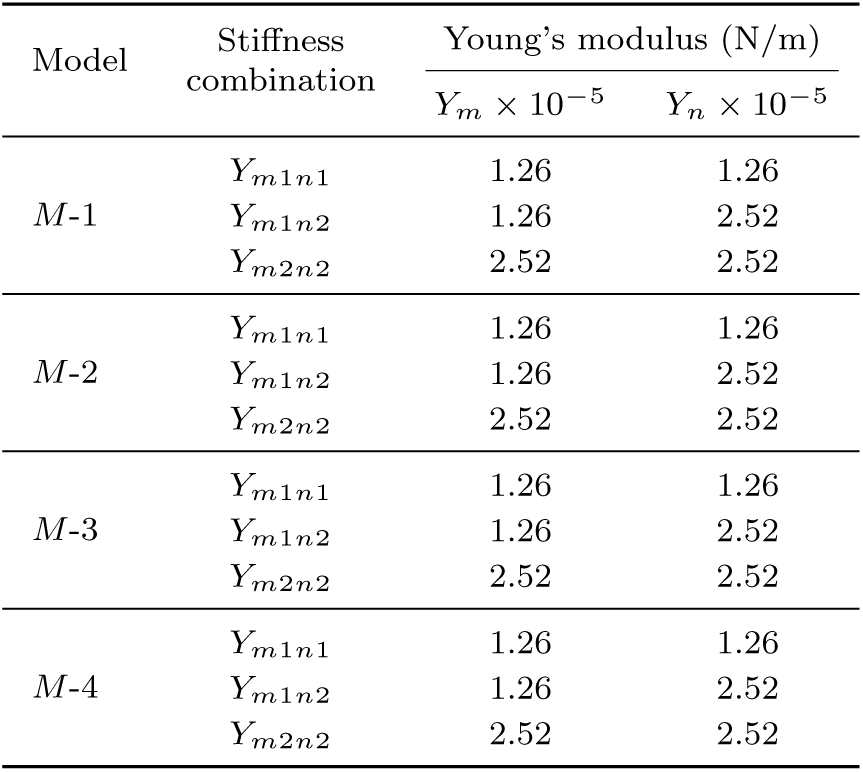
Summary of simulation cases. Each model (*M* −1 to *M* −4) is simulated with three membrane-nucleus stiffness combinations (*Y_m_*_1*n*1_, *Y_m_*_1*n*2_, *Y_m_*_2*n*2_). *Y_m_* and *Y_n_* denote the Young’s moduli of the cell membrane and nucleus, respectively. Δ*t* is the non-dimensional time step size.

| Model | Stiffness combination | Young's modulus (N/m) |  |
| --- | --- | --- | --- |
| | | $Y_m \times 10^{-5}$ | $Y_n \times 10^{-5}$ |
| $M-1$ | $Y_{m1n1}$ | 1.26 | 1.26 |
| | $Y_{m1n2}$ | 1.26 | 2.52 |
| | $Y_{m2n2}$ | 2.52 | 2.52 |
| $M-2$ | $Y_{m1n1}$ | 1.26 | 1.26 |
| | $Y_{m1n2}$ | 1.26 | 2.52 |
| | $Y_{m2n2}$ | 2.52 | 2.52 |
| $M-3$ | $Y_{m1n1}$ | 1.26 | 1.26 |
| | $Y_{m1n2}$ | 1.26 | 2.52 |
| | $Y_{m2n2}$ | 2.52 | 2.52 |
| $M-4$ | $Y_{m1n1}$ | 1.26 | 1.26 |
| | $Y_{m1n2}$ | 1.26 | 2.52 |
| | $Y_{m2n2}$ | 2.52 | 2.52 |

## 3 Results

### 3.1 Cellular Deformation

The shapes of cancer cell observed during simulation at time instants *t*_1_ = 5 ms and *t*_2_ = 10 ms for the four models (Table 4), are shown in Figure 3. Each snapshot in Figure 3 is labeled by its shape class (Table 5). *M* −1 remains streamer-like (*SC*) at both instants for all stiffness combinations, confirming an inherently tethered, highly elongated deformation mode. *M* −2 consistently stays amoeboid-ellipsoidal (*AE*) at *t*_1_ and *t*_2_, indicating stable compact elongation without tether formation. *M* −3 shows the strongest stiffness-driven bifurcation. It is compact-ellipsoidal (*CE*) for all cases at *t*_1_, but by *t*_2_ the softer combinations remain compact (*CE*) or become rounder (*R*), whereas the stiffest case (*Y_m_*_2*n*2_) transitions to a streamer state (*SC*), demonstrating a stiffness-induced switch from compact deformation to tether-dominated elongation. *M* −4 undergoes a clear temporal transition. It is elongated (*EL*) at *t*_1_ for the softer membrane cases and becomes streamer-like (*SC*) for the stiffest configuration, but by *t*_2_ all stiffness combinations converge to the deformed/lobed (*DL*) class, indicating that lobing and folding dominate the loss of compactness rather than continued monotonic stretching. Together, these classifications reinforce that baseline morphology sets the primary deformation mode (persistent *SC* in *M* −1 and persistent *AE* in *M* −2), while stiffness can redirect the deformation pathway in transition-prone geometries (*M* −3 and *M* −4).

**Fig. 3.**
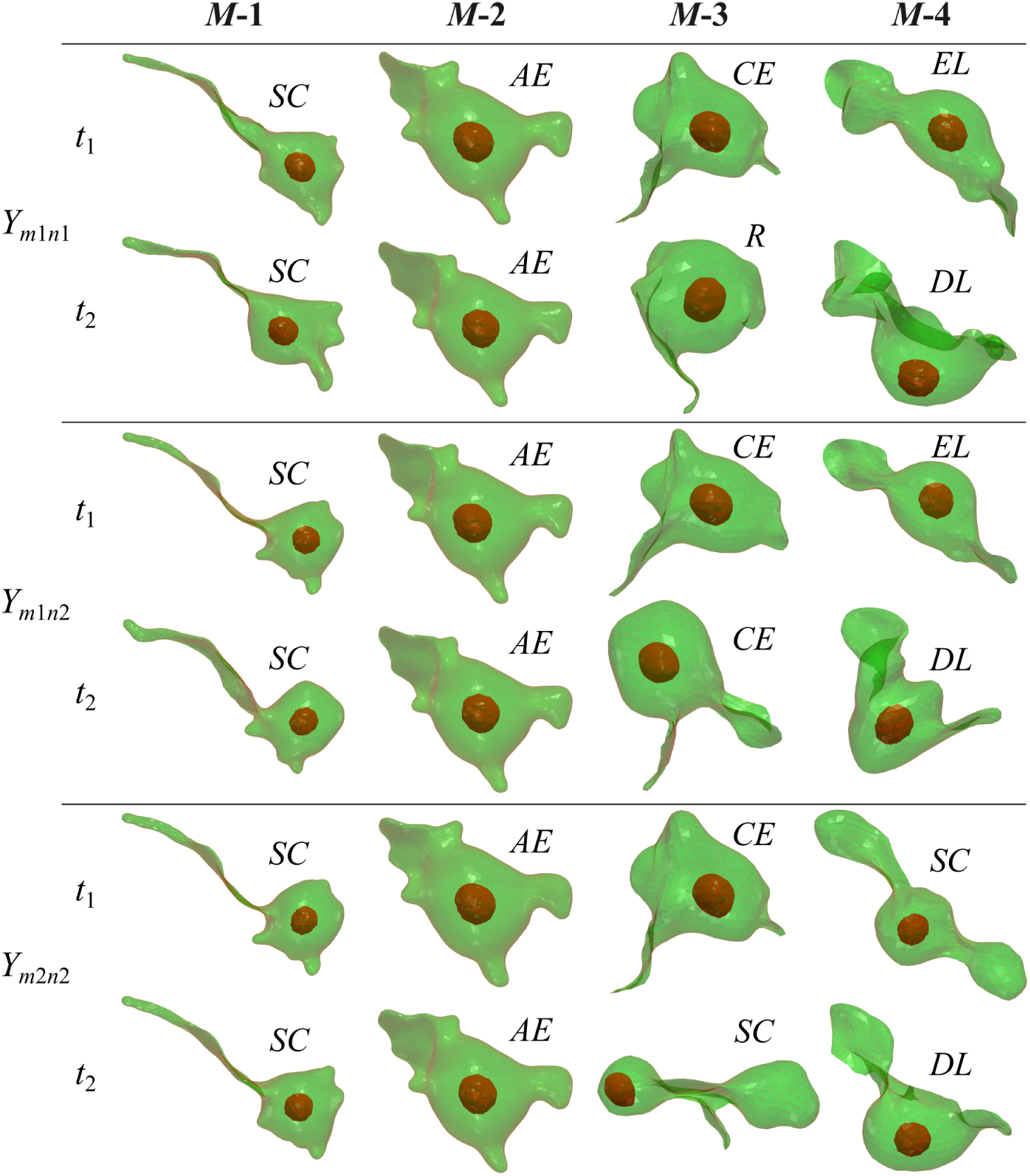
Morphological evolution of the four cancer-cell models (*M* −1, *M* −2, *M* −3, and *M* −4) at two time instants, *t*_1_ = 5 ms and *t*_2_ = 10 ms, for the three stiffness combinations *Y_m_*_1*n*1_, *Y_m_*_1*n*2_, and *Y_m_*_2*n*2_ (Table 4). Each deformed configuration is annotated with its corresponding shape class (*AE*, *R*, *CE*, *EL*, *DL*, and *SC*), based on the sphericity and aspect-ratio (Table 5).

**Table 5.** Classification of deformation-shapes into categories based on the ranges of sphericity (*S*) and aspect ratio (*AR*).

| Class | Label | Criteria in $(S, AR)$ space | Interpretation |
| --- | --- | --- | --- |
| $R$ | Round/compact | $AR \leq 1.8$ and $S \geq 0.77$ | Near-spherical, highest compactness and lowest elongation. |
| $CE$ | Compact-ellipsoidal | $1.8 < AR \leq 3.5$ and $S \geq 0.72$ | Moderately elongated but compact. |
| $AE$ | Amoeboid-ellipsoidal | $3.5 < AR \leq 5.5$ and $S \geq 0.72$ | Relatively higher elongation with high compactness. |
| $DL$ | Deformed/lobed | $AR \leq 5.5$ and $S < 0.72$ | Compactness loss due to lobing/folding, can occur at low-to-moderate $AR$. |
| $EL$ | Elongated | $5.5 < AR \leq 10$ | Strong stretching without extreme tethering. |
| $SC$ | Streamer/comet-like | $AR > 10$ | Tether-dominated extreme elongation. |

Figure 4 quantifies deformation of the four cancer-cell models (Table 4) using the time evolution of aspect ratio (*AR*) and sphericity (*S*). All models respond rapidly within the first 1-2 ms, followed by morphology- and stiffness-dependent evolution. *M* −1 starts highly elongated (*AR >* 10) with *S >* 0.72 (streamer-like, *SC*), then *AR* drops to ∼ 10-12 during *t* ∼ 0.5-1.5 ms placing the cell in *EL*. During *t* ∼ 1.5-6 ms, *AR* remains ≈ 10-13 while *S* decreases from ∼ 0.72 to ∼ 0.62, shifting the morphology toward *DL*. At late times (*t* ∼ 6-10 ms), *Y_m_*_1*n*2_ shows recovery toward *SC* with *AR* ∼ 14-15, whereas *Y_m_*_1*n*1_ and *Y_m_*_2*n*2_ remain mostly in the *EL/SC* range while *S* continues to decline toward ∼ 0.6. Membrane stiffening (*Y_m_*_1*n*2_ to *Y_m_*_2*n*2_) slightly increases *S* and moderates the *AR* evolution, whereas nucleus stiffening (*Y_m_*_1*n*1_ to *Y_m_*_1*n*2_) increases late-time (*t* ≈ 6-10 ms) *AR* variability. *M* −2 begins in the *AE* class (*AE*, *AR* ≈ 5.0-5.3, *S* ≈ 0.81-0.83) and undergoes a brief early compaction. During *t* ∼ 0.2-1.2 ms, *AR* drops to ≈ 2.8-3.4 while *S* remains high (∼ 0.83), placing it in *CE*. It then gradually re-elongates over *t* ∼ 1.2-6 ms (*AR* ∼ 4.2-4.8, *S* ∼ 0.74-0.76), recovering to *AE*. By *t* ∼ 6-10 ms, *AR* plateaus at ∼ 4.6-5.0 for *Y_m_*_1*n*1_ and *Y_m_*_1*n*2_ (slightly lower, ∼ 4.4-4.6, for *Y_m_*_2*n*2_), while *S* approaches ∼ 0.72 for *Y_m_*_1*n*1_ and *Y_m_*_1*n*2_ and remains higher (∼ 0.74-0.75) for *Y_m_*_2*n*2_, thus, *M* −2 remains *AE* with only minor stiffness sensitivity, dominated by membrane effects (lower *AR*, higher *S*). Stronger stiffness dependence emerges in *M* −3 and *M* −4. *M* −3 starts with *CE*, transitions to *AE* during *t* ∼ 1-4 ms (3.5 *< AR* ≤ 5.5), and then to *EL* (5.5 *< AR* ≤ 10). Around *t* ∼ 4.5-6 ms, *AR* ≲ 5.5 with *S <* 0.72 shifts the morphology to *DL*. During *t* ∼ 6-10 ms, only *Y_m_*_2*n*2_ undergoes a sharp rise to *AR >* 10 (transition to *SC*), whereas *Y_m_*_1*n*1_ and *Y_m_*_1*n*2_ remain *DL*. Stiffness effects are negligible up to *t* ≈ 4-5 ms, subsequently, membrane stiffening increases *AR* and decreases *S*, while nucleus stiffening increases both *AR* and *S* over *t* ≈ 5-8 ms before converging. *M* −4 begins elongated (*EL*, *AR* ≈ 8.5, *S* ≈ 0.8) and rapidly becomes streamer-like (*SC*) as *AR >* 10 and *S* drops to ∼ 0.62 during *t* ∼ 0.5-2.5 ms. In the intermediate stage (*t* ∼ 2.5-6 ms), *Y_m_*_1*n*1_ oscillates between *SC* (peaks *AR* ∼ 11) and *EL* (*AR* ∼ 6-8) with *S <* 0.72, *Y_m_*_1*n*2_ shifts to *DL* (*AR* ≲ 5.5, *S* ∼ 0.63-0.66), and *Y_m_*_2*n*2_ reaches the strongest elongation (*AR* ∼ 14-15) before abruptly relaxing from *SC* to *EL* near *t* ∼ 5 ms. At late times (*t* ∼ 6-10 ms), all cases converge to *DL* (*AR* ≲ 5.5, *S <* 0.72). Membrane stiffening produces the strongest early elongation and modifies the relaxation dynamics, but has negligible effect on *S*, whereas nucleus stiffening has little influence on either *AR* or *S*. Overall, membrane stiffness is the dominant regulator of elongation and compactness-particularly for the deformation-prone morphologies (*M* −3 and *M* −4), while nucleus stiffness mainly modulates the timing and amplitude of deformation excursions by constraining interior reconfiguration.

**Fig. 4.**
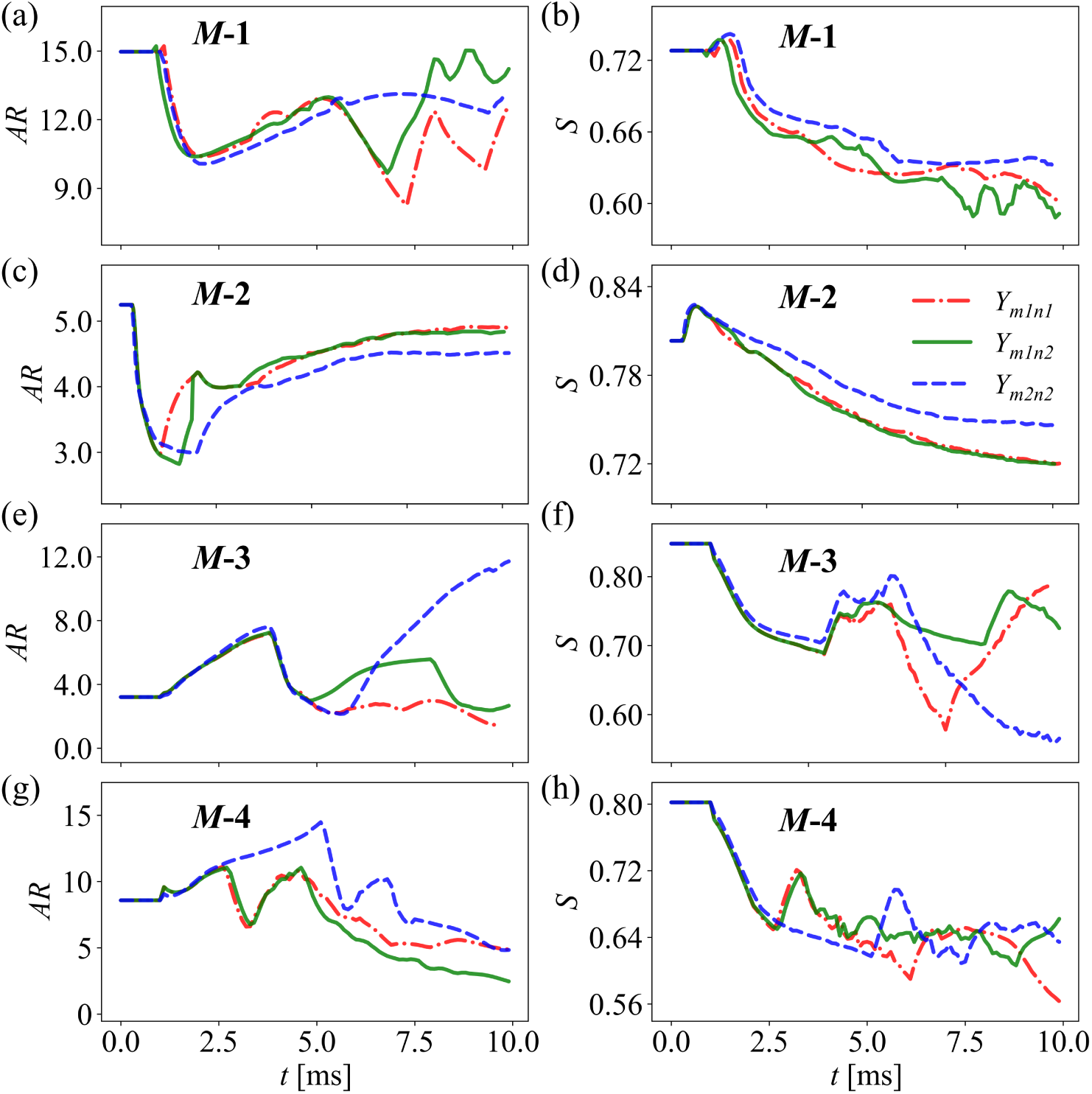
(a, c, e, and g) shows the time evolution of the aspect ratio (*AR*) and (b, d, f, and h) shows sphericity (*S*) for the four cancer-cell models *M* −1, *M* −2, *M* −3, and *M* −4 and three stiffness combinations (*Y_m_*_1*n*1_, *Y_m_*_1*n*2_, and *Y_m_*_2*n*2_).

### 3.2 Near-Field Flow Around Deforming Cancer Cells

Figure 5 compares the extracellular streamlines around the four cancer-cell morphologies (Table 4) at *t*_1_ = 5 ms and *t*_2_ = 10 ms. In *M* −1 the membrane stiffening (comparing *Y_m_*_1*n*2_ and *Y_m_*_2*n*2_) causes the vortices on the top wake region of the cell to combine and form more coherent structure around the top of the cell at *t*_1_. While at *t*_2_ the vortices forming around the cell are washed away due to membrane stiffening. Increasing nuclear stiffness (from *Y_m_*_1*n*1_ to *Y_m_*_1*n*2_) induces cell-shape deformation such that it triggers vortex formation in the wake at *t*_1_, while at *t*_2_, this wake vortex broadens and intensifies. In *M* −2, the shape remains *AE* so the streamlines remain nearly uniform with negligible bending around the cell, indicating that membrane and nuclear stiffening have minimal influence on the surrounding flow field. In *M* −3, during membrane stiffening at *t*_1_ the shape remains *CE* which induces greater streamline deflection around the cell, and by *t*_2_ the shape changes from *CE* to *SC* which shifts the upstream vortex to downstream. During nuclear stiffening at *t*_1_ the shape remains *CE*, causing the streamlines to move closer to the cell, while at *t*_2_ the shape changes from *R* to *CE* so the vortical structures become elongated and advect toward the cell surface. In *M* −4 at *t*_1_ the membrane stiffening leads to washing away of the top and bottom vortices forming in the wake region of the cell as the shape changes from *EL* to *SC*. While at *t*_2_ the shape remains *DL*, so, the vortices are merely stretched and pushed closer to the cell. In contrast, during nuclear stiffening at *t*_1_ the shape remains *EL*, which promotes a reorganization of the vortices around the cell. While at *t*_2_ the shape remains *DL*, causing reorganization of vortices in to more symmetric vortex pattern. Overall, if the membrane stiffening does not change the shape significantly then the vortices are merely stretched and pushed closer to the cell. While a major shape change causes stretching and washing away of the vortices. In contrast, nucleus stiffening mainly reorganizes the vortices when there is not major shape change.

**Fig. 5.**
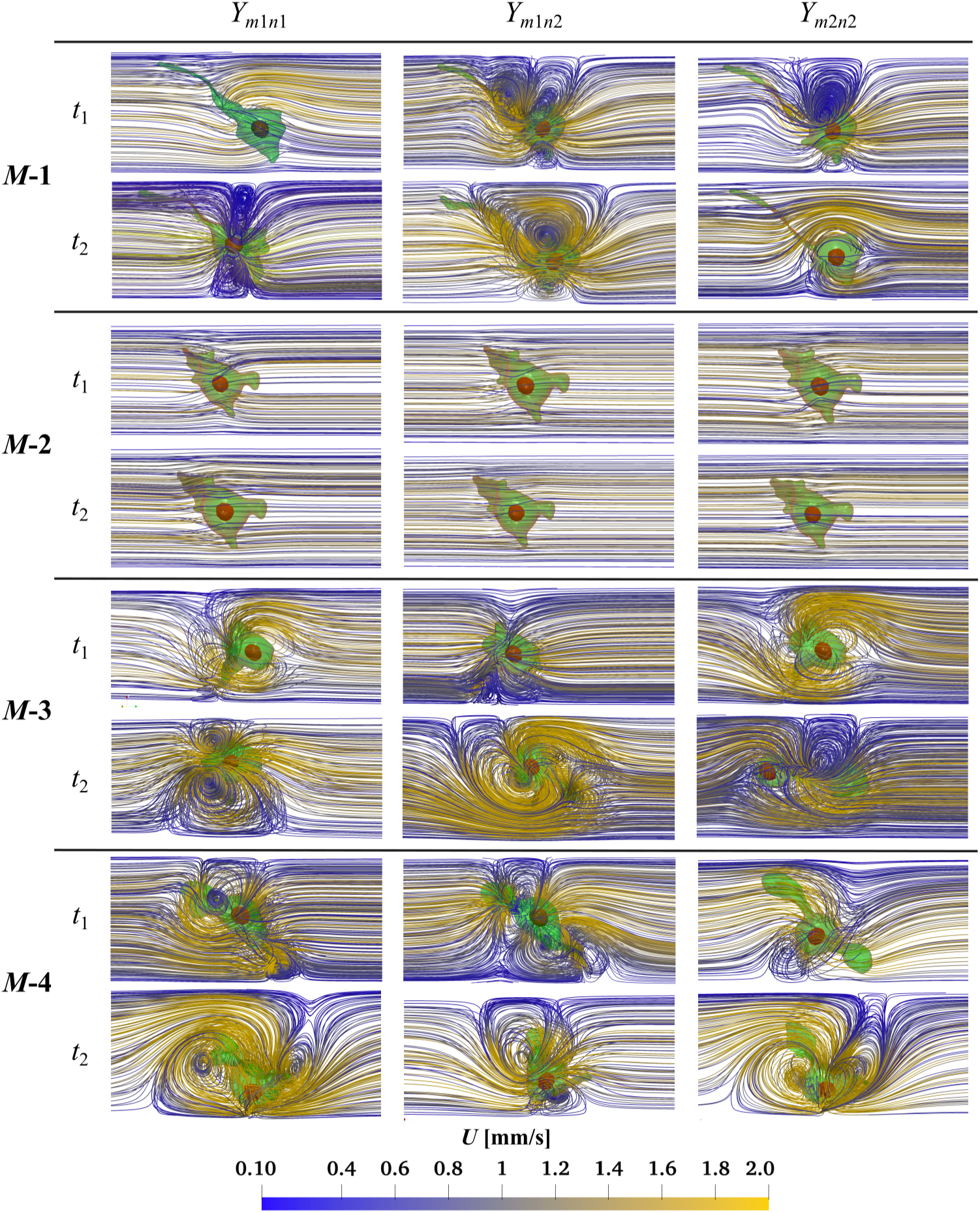
Effect of membrane and nucleus stiffness on the extracellular flow patterns surrounding each of the cancer cell models (*M* −1 to *M* −4) for three stiffness combinations, *Y_m_*_1*n*1_, *Y_m_*_1*n*2_, and *Y_m_*_2*n*2_ (Table 4) at time instants *t*_1_ = 5 ms and *t*_2_ = 10 ms for three stiffness combinations. Across all morphologies, increasing stiffness makes the cell behave more obstacle-like, leading to more localized recirculation, although the strength of this response depends primarily on the underlying morphology.

These stiffness-dependent changes in the extracellular flow directly influence the surface loading on the membrane through pressure and shear stresses. Therefore, Figure 6 reports the traction magnitude, defined as the combined contribution of pressure and viscous shear acting on the membrane surface, for all the four morphologies (Table 4) at *t*_1_ and *t*_2_. The traction distribution again reflects a strong dependence on baseline shape. The compact morphology *M* −2 exhibits the smoothest and most symmetric loading, with moderate traction levels mainly concentrated near the front and rear regions and only weak sensitivity to stiffness. In contrast, the deformation-prone morphologies (*M* −1, *M* −3, and *M* −4) show pronounced traction localization near geometric constrictions, high-curvature regions, and neck/tether-like features, and these hot spots intensify and reorganize from *t*_1_ to *t*_2_ as deformation progresses. Membrane stiffening sharpens the traction gradients and concentrates the maxima into smaller, more intense patches by reducing surface compliance. Nucleus stiffening limits internal shape changes, which redistributes the traction field and typically shifts the highest traction toward the outer surface. As a result, traction localization remains weak for *M* −2, but becomes prominent for *M* −1, *M* −3, and especially *M* −4, where stiffness-dependent deformation strongly rearranges the dominant loading zones between *t*_1_ and *t*_2_.

**Fig. 6.**
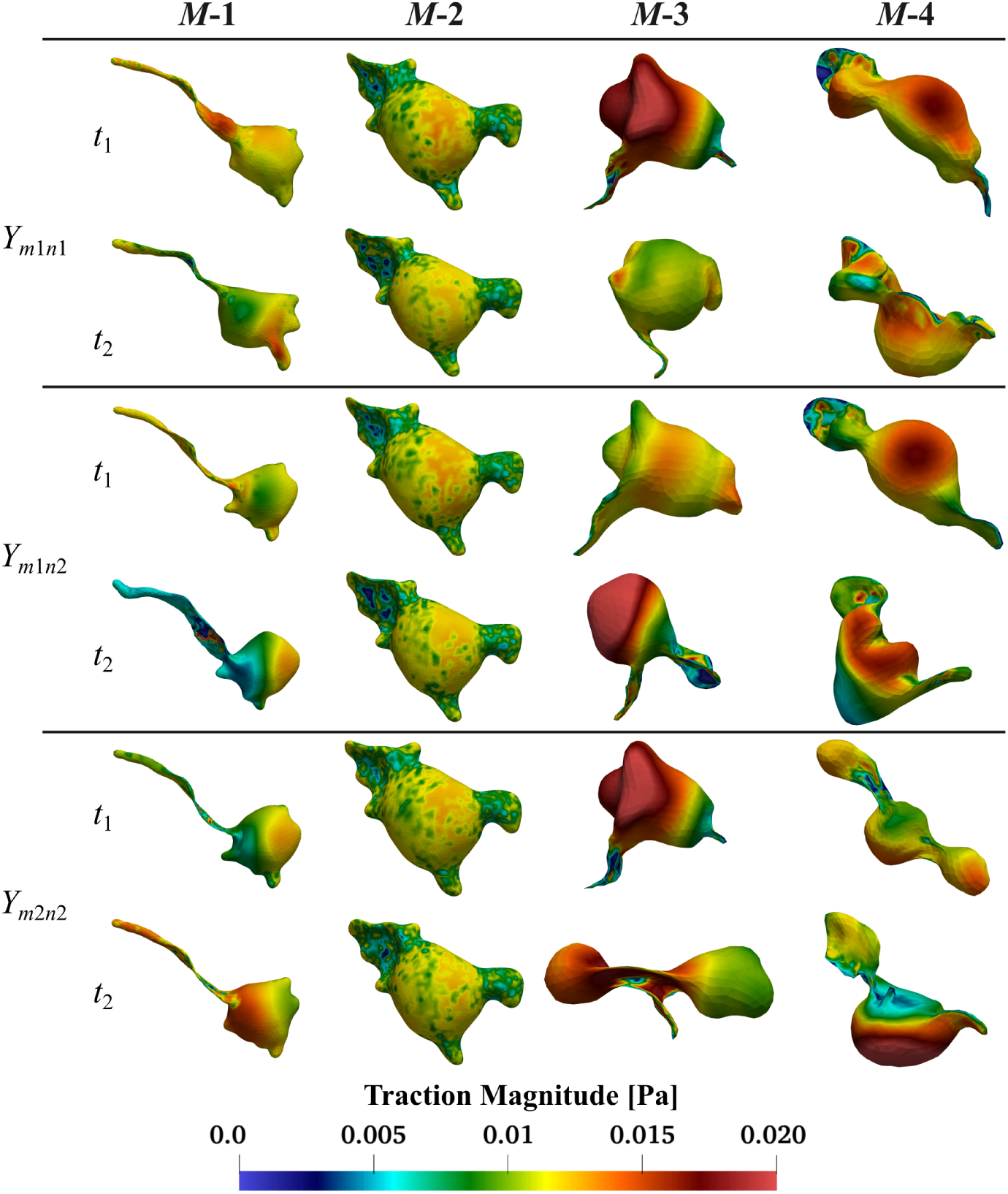
Spatial distribution of membrane traction magnitude on the four cancer-cell morphologies (*M* −1, *M* −2, *M* −3, and *M* −4) at two representative instants (*t*_1_ = 5 ms and *t*_2_ = 10 ms) for the three stiffness combinations (*Y_m_*_1*n*1_, *Y_m_*_1*n*2_, and *Y_m_*_2*n*2_) (Table 4). The traction field highlights where the fluid-structure interaction loads concentrate on the evolving surface and therefore provides a spatial complement to the integrated force histories.

### 3.3 Total Hydrodynamic Force on the Membrane

Figure 7 shows the variation of net hydrodynamic force on the membrane for cancer cell models *M* −1, *M* −2, *M* −3, and *M* −4 for three stiffness combinations *Y_m_*_1*n*1_, *Y_m_*_1*n*1_, and *Y_m_*_1*n*1_ (Table 4). Across all morphologies, the total membrane force shows the same start-up response. It rises sharply as the flow is established within *t* ≈ 0-0.5 ms. It then relaxes toward a quasi-steady level. The differences between the stiffness cases become clearer after *t* ≈ 1 ms. These differences arise from changes driven by deformation in the effective blockage and wake structure. For *M* −1, the force stabilizes quickly. It settles near *F* ≈ 18-20 pN and remains slightly unsteady through *t* = 10 ms. The three stiffness curves remain close. They differ slightly during *t* ≈ 6-10 ms. For *M* −2, the response is the most stable. After the start-up transient, the curves nearly overlap. The force remains almost constant at *F* ≈ 14-16 pN from *t* ≈ 1 ms to *t* = 10 ms. This indicates a weak stiffness sensitivity for this compact morphology. For *M* −3, all cases relax to a similar level after starting. The force is typically *F* ≈ 16-18 pN over *t* ≈ 1-5 ms. Stiffness controls the transient excursions afterward. The most compliant case *Y_m_*_1*n*1_ shows a pronounced overshoot. It reaches ∼ 30 pN around *t* ≈ 6-7 ms. Then it drops to a lower level by *t* ≈ 8-10 ms. The stiffer cases suppress this peak and evolve more smoothly. For *M* −4, the force remains strongly time-dependent. After the initial rise, it continues to increase and becomes more unsteady during *t* ≈ 2-10 ms. The highest values occur near *t* = 10 ms. The force reached about 25-27 pN for the stiffest cases. In general, compact morphologies produce lower and steadier forces. They show minimal stiffness dependence as seen in *M* −2. Deformation-prone morphologies show stronger unsteadiness and clearer modulation of stiffness. This behavior is most evident in *M* −3 and *M* −4.

**Fig. 7.**
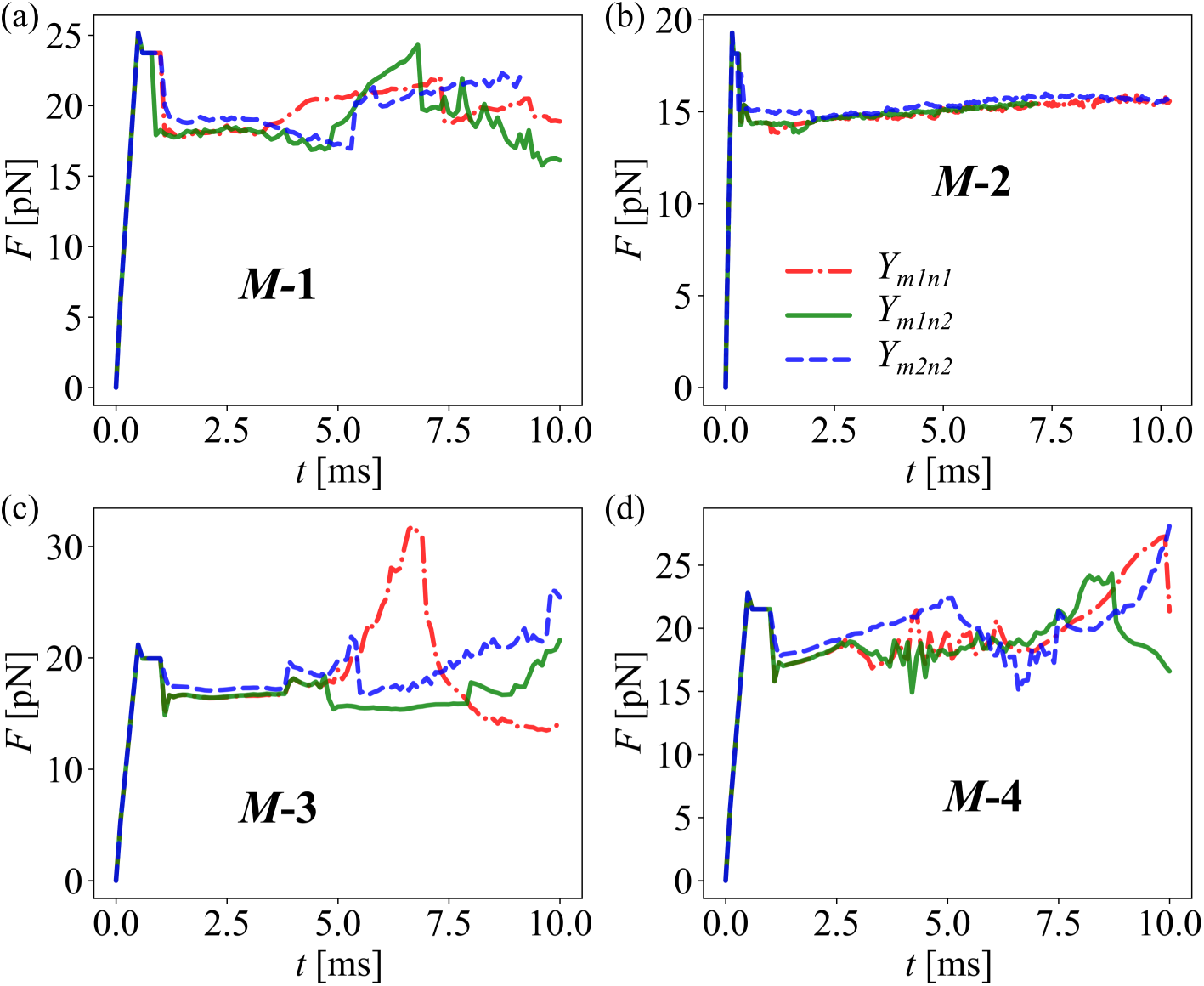
Temporal profile of the total hydrodynamic force magnitude *F* acting on the cancer cell membrane for the four cell morphologies *M* −1,*M* −2,*M* −3, and *M* −4 under uniform flow. Results are shown for three membrane and nucleus stiffness combinations, *Y_m_*_1*n*1_, *Y_m_*_1*n*2_, and *Y_m_*_2*n*2_ (See Table 4).

In Fig. 8, the transverse velocity and vorticity fields around *M* −1 show the symmetry breaking about *xx*- and *yy*-axes that drives its lateral migration. The velocity contour on the first plane at *t*_1_ for stiffness *Y_m_*_1*n*1_ shows that velocity distribution is nearly symmetric about *xx* while significant asymmetry can be observed about *yy*. Similar trends are observed in other two planes. Higher asymmetry about *yy* will cause higher shear and pressure force imbalance in *x*-direction as compared to *y*-direction, which will cause the dominant displacement in *x*-direction as compared to *y*-direction. Membrane stiffening (*Y_m_*_1*n*2_ to *Y_m_*_2*n*2_) and nucleus stiffening (*Y_m_*_1*n*1_ to *Y_m_*_1*n*2_) only modulate the intensity of the contours without effecting the trend significantly. Similar trend can be observed at *t*_2_ with variation in velocity intensity. Consistent with the velocity contour plots vorticity contour plots also show greater symmetry across *xx*-axis as compared to *yy*-axis. Therefore, they support the previous finding that displacement in *x*-direction will be greater than *y*-direction due to higher unbalanced force about *yy*-axis as compared to *xx*-axis. The weaker asymmetry about the *xx*-axis leaves only a small residual imbalance in the *y*-direction and therefore limited migration in that direction.

**Fig. 8.**
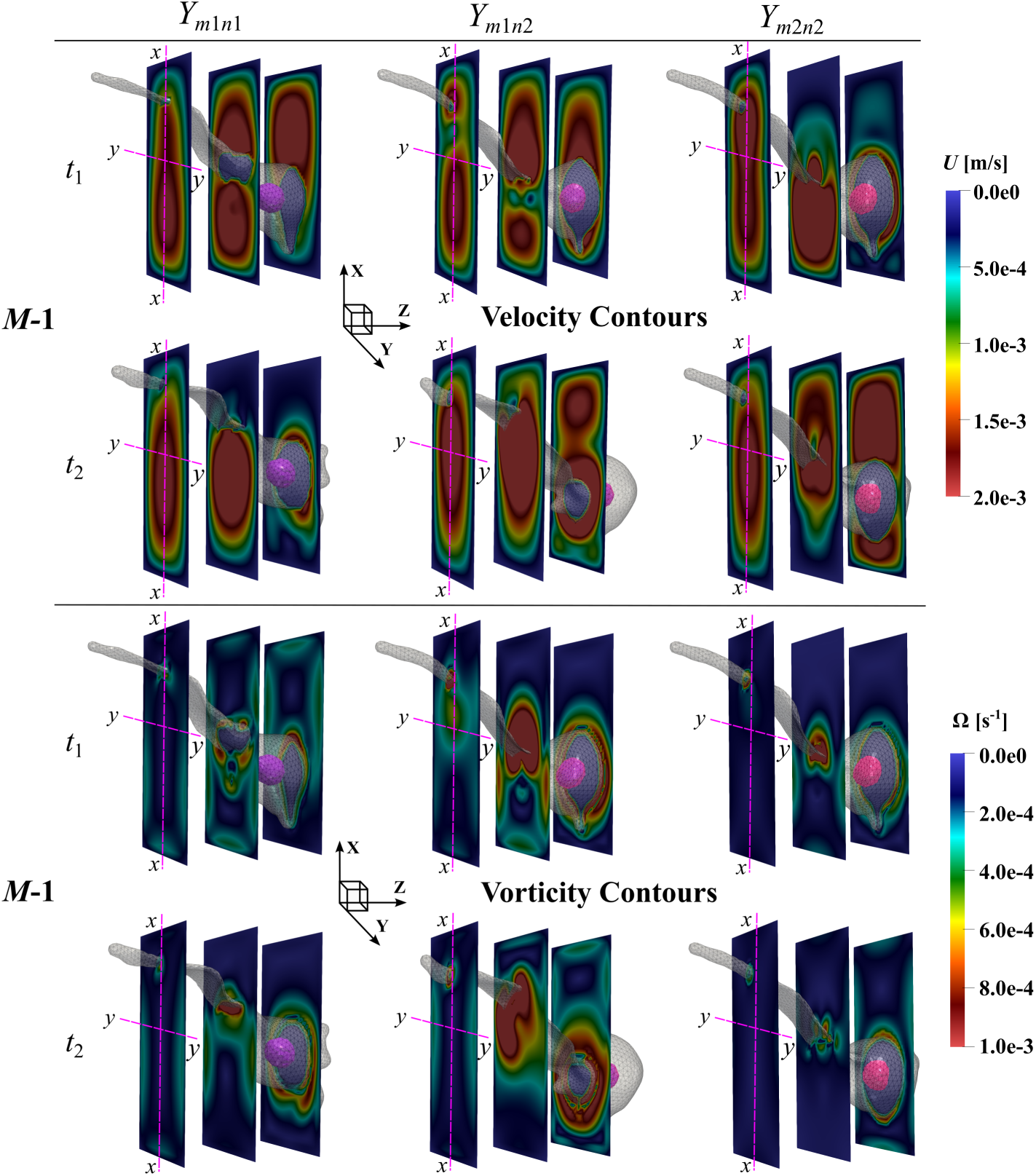
Velocity magnitude (*U*) and vorticity magnitude (Ω) contours on three transverse planes (normal to the flow direction *z*) for cancer-cell model *M* −1 at two representative time instants, *t*_1_ = 5 ms and *t*_2_ = 10 ms. Results are shown for the three stiffness combinations *Y_m_*_1*n*1_, *Y_m_*_1*n*2_, and *Y_m_*_2*n*2_ (Table 4). *xx*- and *yy*-axes on the first plane are used to show the asymmetry in the velocity and vorticity contours which causes drift in transverse direction.

## 4 Discussion

A broad range of computational frameworks has been developed to study cancer-cell transport in flow, aiming to identify the mechanical factors that regulate key steps of the metastatic cascade [42]. Particle-based methods [43], particularly dissipative particle dynamics (DPD), remain widely used for cell-scale hydrodynamics and adhesion problems [44]. Recent DPD studies examined tumor-cell trajectories in bifurcations [45] and adhesion in curved microvessels [26]. However, many prior simulations assume idealized initial morphologies (e.g., spherical/capsule-like cells), which does not fully represent in situ morphological heterogeneity [22] and may under-estimate morphology-driven differences in wake asymmetry, traction localization, and cross-stream forcing. In this study, we resolve deformation and membrane loading, for cancer cells with realistic membrane morphologies while systematically varying membrane and nuclear stiffness.

The current study shows that Cancer-cell transport in confined flow reflects a coupled morphology-mechanics-hydrodynamics response in which baseline geometry sets the dominant deformation pathway, while membrane and nuclear stiffness tune its intensity, timing, and recovery (Figs. 3, 4). Morphology and stiffness dependent changes in shapes indices (*AR*, *S*, Figure 4) and total hydrodynamic force (*F*, Figure 7) were observed only after ∼ 1-2 ms after the start of flow.

Cancer cell *M* −1, *AR* partially recovers within *t* ≈ 4–6 ms (Figure 4a,b). While *S* converses from ∼ 0.72 towards ∼ 0.60–0.64 without showing any recovery. *M* −2 exhibits the clearest elastic recovery in geometry (Figure 4b,c). *AR* increases steadily and approaches a stable plateau by *t* ≈ 6–10 ms, reflecting a smooth relaxation toward a steady elongated shape without unstable tethering. In contrast, *S* decreases gradually and does not show a strong rebound. For *M* −3, after the initial drop in *S* and rise in *AR*, the cell undergoes a clear reorganization around *t* ≈ 4–6 ms, where *AR* temporarily decreases and *S* temporarily increases (partial recovery in elongation and compactness, Figure 4e,f). After this point, recovery becomes stiffness-controlled, the softer cases show partial restoration of elongation and compactness, whereas the stiffest case sharply decline in *S* and a sustained rise in *AR*, showing no recovery. *M* −4 displays repeated deformation–relaxation cycles (Figure 4g,h). *AR* shows strong oscillations, indicating alternating stretching and partial relaxation events, while *S* drops quickly to ∼ 0.63–0.66 by *t* ≈ 2–3 ms and then fluctuates around this lower level. Overall, the compact shapes (*M* −2, *M* −3) show significant and consistent shape recovery as compared to the elongated shapes (*M* −1, *M* −4). The previous literature also report similar finding, Kim *et al.* [46] used mechano-node-pore sensing method which measures cell deformation and recovery time after deformation shows that malignant cells have distinct mechanical recovery phenotypes compared to non-malignant cells. While the study of deformation of cencer cells when they pass through a constricted channel showed recovery after peak deformation [47].

Near-field flow responses are consistent with this shape-governed framework as shown in Figure 5. When stiffness does not induce a major shape-class change (Table 5), vortical structures tend to stretch and move closer to the cell surface, when a major transition occurs (*CE* to *SC* in *M* −3 or *EL* to *SC* in *M* −4), vortices reorganize strongly or are washed away, reflecting altered separation and wake dynamics (Figure 5). This coupling is further reflected in traction distributions: *M* −2 maintains the smoothest and most symmetric loading with weak stiffness sensitivity, whereas *M* −1, *M* −3, and *M* - 4 develop pronounced traction localization near constrictions, high-curvature regions, and tether-like features (Figure 6). Therefore, membrane stiffening reduces surface compliance and concentrates the fluid-induced loading into smaller, more intense traction hot spots, consistent with experimental observations that increased membrane stiffness (via cholesterol depletion) elevates resistance to deformation and strengthens local mechanical gradients [48]. In contrast, nucleus stiffening limits internal rearrangement and nuclear deformation, which redistributes load pathways through the cytoskeleton and shifts where peak stresses accumulate on the cell surface [49].

The integrated force histories (Figure 7) provide a complementary view of deformation mechanisms and clarify when stiffness effects become dynamically relevant. All cases show a start-up rise within *t* ≈ 0-0.5 ms followed by relaxation (Figure 7). Differences become clearer after *t* ≈ 1 ms when shape evolution begins to alter effective blockage and wake structure. The compact morphology *M* −2 exhibits the lowest unsteadiness and the weakest stiffness sensitivity (Figure 7b), consistent with limited wake reorganization (Figure 5). In contrast, *M* −3 and *M* −4 show the clearest stiffness modulation of force unsteadiness and amplitude (Figure 7c, d). In *M* −3, the large overshoot in the compliant case around *t* ≈ 6-7 ms indicates that transient shape reorganization (Figure 3) can amplify hydrodynamic loading. The suppression of this overshoot in stiffer configurations supports the interpretation that stiffness constrains the deformation pathway and limits abrupt changes in effective blockage [50]. In *M* −4, the progressive force growth and increasing unsteadiness over *t* ≈ 2-10 ms reflect persistent asymmetric deformation (Figure 3) and continual wake reorganization (Figure 5), which is consistent with the heterogeneous traction patterns (Figure 6) and the transition to lobed configurations by *t*_2_ (Figure 3).

Finally, the transverse velocity and vorticity fields (Figure 8) rationalize the migration tendency of *M* −1 and clarify why drift is expected to be stronger in the *x*-direction than in the *y*-direction. The contours show a more pronounced asymmetry about the *yy*-axis than about the *xx*-axis at both *t*_1_ and *t*_2_. This implies that the velocity deficit and the associated shear-layer roll-up are biased laterally in a way that produces a larger imbalance of pressure and shear stresses in the *x*-direction [51]. The corresponding imbalance in the *y*-direction is weaker because the asymmetry about the *xx*-axis is smaller. Membrane and nucleus stiffening primarily modulate the intensity of these features by sharpening gradients and concentrating vorticity. They do not change the dominant symmetry-breaking direction. This supports the interpretation that morphology-driven wake bias sets the migration directionality [52].

Overall, these findings suggest that mechanical phenotype should be interpreted through deformation pathways rather than deformation magnitude alone. Membrane stiffness plays the leading role by selecting whether the cell remains in compact elongation, develops tethered streamer-like states, or transitions toward lobed configurations. Nucleus stiffness plays a secondary role by constraining interior rearrangement and shifting where deformation and loading localize. This coupling between morphology and stiffness is the key reason that transition-prone geometries exhibit the strongest sensitivity in near-field flow, traction localization, and force unsteadiness. Future work that adds adhesive kinetics, cell-cell interactions, and spatially varying shear will help translate these mechanistic links into predictions for vascular arrest and extravasation [53] under physiologically realistic conditions.

## 5 Conclusion

This work presents the first simulation of cancer-cell dynamics using a high-fidelity hybrid continuum-particle framework [54–56]. Cancer-cell geometries reconstructed from microscopic imaging data were used to quantify how realistic morphology, together with membrane and nucleus stiffness, governs transient deformation and migration in shear-driven microchannel flows. The approach also resolves extracellular flow structures while providing surface-resolved estimates of hydrodynamic loading.

Our simulations show that cell morphology controls extracellular hydrodynamics and traction, which consequently leads to stiffness modulated cell deformation. Cancer-cell response to shear flow is primarily governed by baseline shape. Within that morphology-driven framework, membrane stiffness controls elongation and the localization of traction hot spots on the surface, whereas nucleus stiffness further modulates the outcome by limiting internal rearrangement and steering the deformation pathway. Compact morphologies produce nearly symmetric fluid flow velocity deficits in the cell wake, leading to negligible cross-stream drift. In contrast, elongated and lobed morphologies generate laterally displaced wakes and localized vorticity hot spots, which maintained a transverse hydrodynamic imbalance and produced measurable lateral migration.

Our results therefore emphasize the stiffness- and shape-dependent traction distributions provide mechanically consistent precursors which might promotes cell migration toward blood vessel walls [26]. Elongated/streamer-prone shapes more readily sustained or transitioned to highly elongated states when the nucleus was stiff. The compact morphologies showed limited elongation even at high nuclear stiffness. Our future work will include the simulation of receptorligand adhesion, blood-cell interactions, and collective effects to better predict vascular arrest in realistic microvascular flows. [26] under physiological conditions.

## Acknowledgements.

This work is supported by the NSF grant number 1946202 ND-ACES and a start-up package of Trung Le from North Dakota State University. The authors acknowledge the use of computational resources at the Center for Computationally Assisted Science and Technology CCAST-NDSU, which is supported by the NSF MRI 2019077. The authors also received allocation CTS200012 from the Extreme Science and Engineering Discovery Environment (XSEDE). We acknowledge the financial support of NIH-2P20GM103442-19A1 to train undergraduate students in Biomedical Engineering.

## 7 Contributions

Conceptualization: Trung Le, Amanda Haage; Methodology: Amanda Haage, Lahcen Akerkouch, Trung Le; Formal analysis and investigation: Meraj Ahmed, Lahcen Akerouch, Aaron Vanyo; Writing - original draft preparation: Meraj Ahmed, Lahcen Akerkouch, Trung Le; Writing - review and editing: Meraj Ahmed, Trung Le, Amanda Haage; Funding acquisition: Trung Le, Amanda Haage; Resources: Trung Le, Amanda Haage; Supervision: Trung Le, Amanda Haage.

## 8 Data Availability

Data sets generated during the current study are available from the corresponding author on reasonable request.

